# Genetic risk for autism spectrum disorders and neuropsychiatric variation in the general population

**DOI:** 10.1101/027771

**Authors:** Elise B. Robinson, Beate St. Pourcain, Verneri Anttila, Jack Kosmicki, Brendan Bulik-Sullivan, Jakob Grove, Julian Maller, Kaitlin E. Samocha, Stephan Sanders, Stephan Ripke, Joanna Martin, Mads V. Hollegaard, Thomas Werge, David M. Hougaard, iPSYCH-SSI-Broad Autism Group, Benjamin M. Neale, David M. Evans, David Skuse, Preben Bo Mortensen, Anders D. Børglum, Angelica Ronald, George Davey Smith, Mark J. Daly

## Abstract

Almost all genetic risk factors for autism spectrum disorders (ASDs) can be found in the general population, but the effects of that risk are unclear in people not ascertained for neuropsychiatric symptoms. Using several large ASD consortia and population based resources, we find genetic links between ASDs and typical variation in social behavior and adaptive functioning. This finding is evidenced through both inherited and *de novo* variation, indicating that multiple types of genetic risk for ASDs influence a continuum of behavioral and developmental traits, the severe tail of which can result in an ASD or other neuropsychiatric disorder diagnosis. A continuum model should inform the design and interpretation of studies of neuropsychiatric disease biology.

---

Autism spectrum disorders (ASDs) are a group of neuropsychiatric conditions defined through deficits in social communication, as well as restricted and repetitive interests. Consistent with traditional approaches to psychiatric phenotypes, most genetic studies of ASDs compare cases to controls in order to identify risk-associated variation. This approach has been highly productive—recent studies have linked common polygenic as well as *de novo* and inherited rare variation to ASD risk ^2,3^. Common genotyped SNPs are estimated to account for at least 20% of ASD liability ^2,4,5^. Contributing *de novo* variants are found in 10-20% of cases but *de novo* mutations collectively explain less than 5% of overall ASD liability ^2,6,7^.

Almost all genetic risk factors for ASDs can be found in unaffected individuals. For example, most people who carry a 16p11.2 deletion, the most common large mutational risk factor for ASDs, do not meet criteria for an ASD diagnosis^8^. Across healthy populations there is also significant variability in capacity for social interaction and social communication ^9^. While such phenotypic variation is well established, the genetic relationship between neuropsychiatric disorders and typical social and behavioral variation remains unclear. From the first published descriptions of ASDs, clinical and epidemiologic reports have commonly noted subthreshold traits of autism in the family members of many diagnosed individuals ^10,11^. Twin and family studies have suggested that those similarities are at least in part inherited ^12-14^, but a correlation between traits and the diagnosis has yet to be identified using genetic data.

We examined the association between genetic risk for ASDs and social and behavioral variation in the general population, and examined the model through which genetic risk for ASDs is conferred. Traditional categorical psychiatric diagnoses (e.g., yes/no ASD) ignore the possibility of intermediate outcomes, long known to be relevant to phenotypes like intellectual disability and IQ that are more easily quantified. Several studies have now associated neuropsychiatric disease associated CNVs with cognitive or educational differences in the general population ^15,16^. *De novo* deletions at 16p11.2 were recently reported to confer a quantitative effect on intelligence (an average two standard deviation reduction from mean IQ of the parents), rather than creating risk for categorical yes/no intellectual disability^17^. The extent to which such patterns extend to social and behavioral traits is unknown and could substantially influence: 1) the design and interpretation of biological studies of ASDs and other severe mental illnesses, as well as 2) the designation of therapeutic treatment thresholds. We aimed to resolve this question using multiple categories of genetic variation that create risk for ASDs—common, inherited alleles as well as rare, *de novo* variants.

## The genetic correlation between ASDs and typical variation in social and communication ability

Like nearly all common diseases, common variant risk for ASDs is distributed across the genome, with many thousands of contributing loci of small effect ^2,4^. The cumulative contribution of common SNPs to ASD risk (SNP heritability^18^) has been estimated using several methods, most recently and efficiently via LD score regression. LD score regression makes use of GWAS summary statistics to estimate SNP heritability ^5^. The method can also be used to estimate the correlation between common variant influences on two phenotypes (genetic correlation) ^1^. As LD score correlation requires only GWAS summary statistics, genetic correlations can be estimated between distinct data sets and cohorts.

We used three data sets to examine the common variant association between ASDs and social and communication difficulties in the general population (Table S1). First, traits of social and communication impairment were measured using the Social and Communication Disorders Checklist (SCDC) in the Avon Longitudinal Study of Parents and Children (ALSPAC), a general population cohort born in 1991-1992 in Bristol, United Kingdom ^19,20^. The SCDC is a parent-rated quantitative measure of social communication impairment that is continuously distributed and has been extensively studied ^21-23^. There is substantial trait overlap between the SCDC and canonical ASD symptomology (e.g. “Not aware of other people’s feelings”) and children with ASDs have very high average scores (indicating many difficulties) on the SCDC ^21^. The measure does not include items on restricted and repetitive interests. For the purposes of this project, we used summary statistics from a published GWAS of the SCDC, administered when the children were 8 years old (n=5,628) ^23^. The SNP heritability of the SCDC was 0.17 (s.e.=0.09) using LD score regression, similar to the estimate derived from residual maximum likelihood using the software package GCTA (h2g=0.24, s.e.=0.07; n=5,204) ^23^.

We correlated the genetic influences on the SCDC with those on diagnosed ASDs using ASD data from two large consortium efforts. The Psychiatric Genomics Consortium autism group (PGC-ASD) has completed a GWAS of 5,305 ASD cases and 5,305 pseudocontrols constructed from untransmitted parental chromosomes (see Online Methods). Summary statistics from that GWAS are publicly available through the PGC website (URLs: PGC). As a replication set, we recently completed an independent ASD case-control GWAS with 7,783 ASD cases and 11,359 controls from the Danish iPSYCH project (iPSYCH-ASD; Online Methods). Using LD score regression, we estimated that the liability-scale SNP heritability of PGC-ASD was 0.23 (s.e.=0.03; assumed population prevalence 1%), suggesting that approximately one quarter of ASD liability reflects common genotyped variation. The estimated liability-scale SNP heritability of iPSYCH-ASDs was 0.14 (se=0.03; assumed population prevalence 1%). The genetic correlation between PGC-ASD and iPSYCH-ASD was 0.74(se=0.07; p<1e-20), indicating similar common, polygenic architecture between ASDs diagnosed primarily in the United States and Denmark.

The estimated genetic correlations between the SCDC in ALSPAC and the two ASD GWAS data sets are shown in Figure 1. The estimated genetic correlation between PGC-ASD and the SCDC was 0.27 (s.e.= 0.13, p=0.006), suggesting that approximately one quarter of the genetic influences on ASDs share that influence with the SCDC. The estimated genetic correlation between iPSYCH-ASD and the SCDC was very similar (r_g_=0.29; s.e.=0.09; p=0.001), evidencing substantial and replicable etiologic overlap between ASDs and typical variation in social and communication ability in childhood. The estimated genetic correlations between ASDs and the SCDC exceed those previously estimated between PGC ASD and each of PGC schizophrenia, PGC bipolar disorder, and PGC major depressive disorder as shown in Figure 1. This suggests that ASDs are at least as strongly associated with social and communication trait variation in the population as they are several other categorically diagnosed psychiatric disorders.

**Figure 1.**
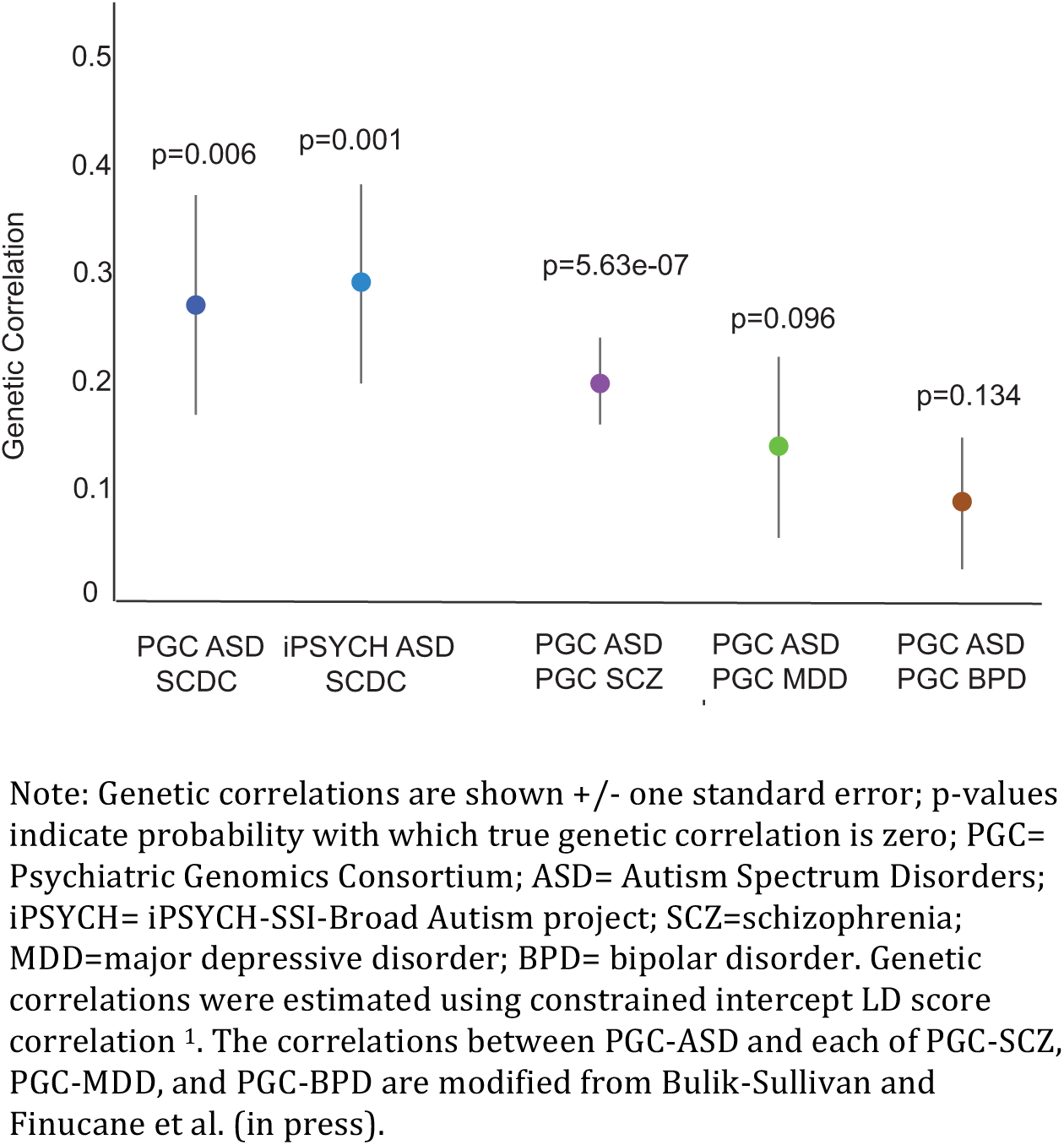
The genetic correlation between ASDs and social and communication difficulties in the general population

We next asked whether the correlation between ASDs and the SCDC might be confounded by a negative association between common variant ASD risk and full scale IQ. In fact, a positive genetic association between PGC-ASD and IQ/educational attainment in the general population has been reported several times ^1,24,25^. One of those studies used IQ summary statistics from a child IQ GWAS (mean age 9 years; Benyamin 2014), from which ALSPAC was the largest contributing dataset (Bulik-Sullivan and Finucane, in press; PGC-ASD to IQ r_g_=0.40, s.e.=0.09, p=5.27e-06). We examined the association between the same child IQ GWAS and iPSYCH-ASD. The positive genetic correlation did not replicate (r_g_= 0.08; se= 0.07; p=0.23), but suggests that the common polygenic relationship between ASDs and IQ in the general population is consistently non-negative. This is particularly intriguing given the strong and consistently negative association between *de novo* variant risk for ASDs and intellectual functioning ^7,26,27^. Genetic correlations from GWAS data are sensitive to case and control selection, for example ascertainment of case families that might be, on average, of higher socioeconomic status than controls. The PGC ASD summary statistics are immune to this sort of confounding, because they were generated using a case/pseudocontrol study design (see Online Methods). Cases and pseudocontrols are matched for genetic load for all phenotypes except those that are genetically correlated with case/control status. This holds irrespective of biases that may exist with respect to the detection and ascertainment of cases and families. The PGC-ASD dataset is therefore particularly well suited for examining the association between common variant risk for ASDs and cognition.

## The continuum of behavioral and developmental outcomes associated with de novo variation

We next aimed to examine whether *de novo* variant data similarly gives evidence of a genetic relationship between ASDs and a continuum of behavioral outcomes in the population. The Simons Simplex Collection (SSC) is a sample of over 2,800 individuals with ASDs and their nuclear family members, with extensive data on unaffected siblings ^28^. To our knowledge, the SSC unaffected siblings are currently the only deeply phenotyped control sample with *de novo* variant information available. The Vineland Scales of Adaptive Behavior (Vineland) was used in the sequenced SSC cases (n=2,497) and sibling controls (n=1,861). The Vineland captures parent-rated variation in social, communication, and daily living skills, and normalizes those abilities to a mean of 100 and standard deviation of 15 in the population, corrected for age ^29^. On average, individuals with ASDs have mean Vineland scores approximately two standard deviations below the mean (SSC case mean = 73.3; s.d. = 12.17), consistent with the social and communication impairments definitional to the diagnosis. The distribution of the Vineland overlaps between cases and controls, as shown in Figure 2a.

**Figure 2a.**
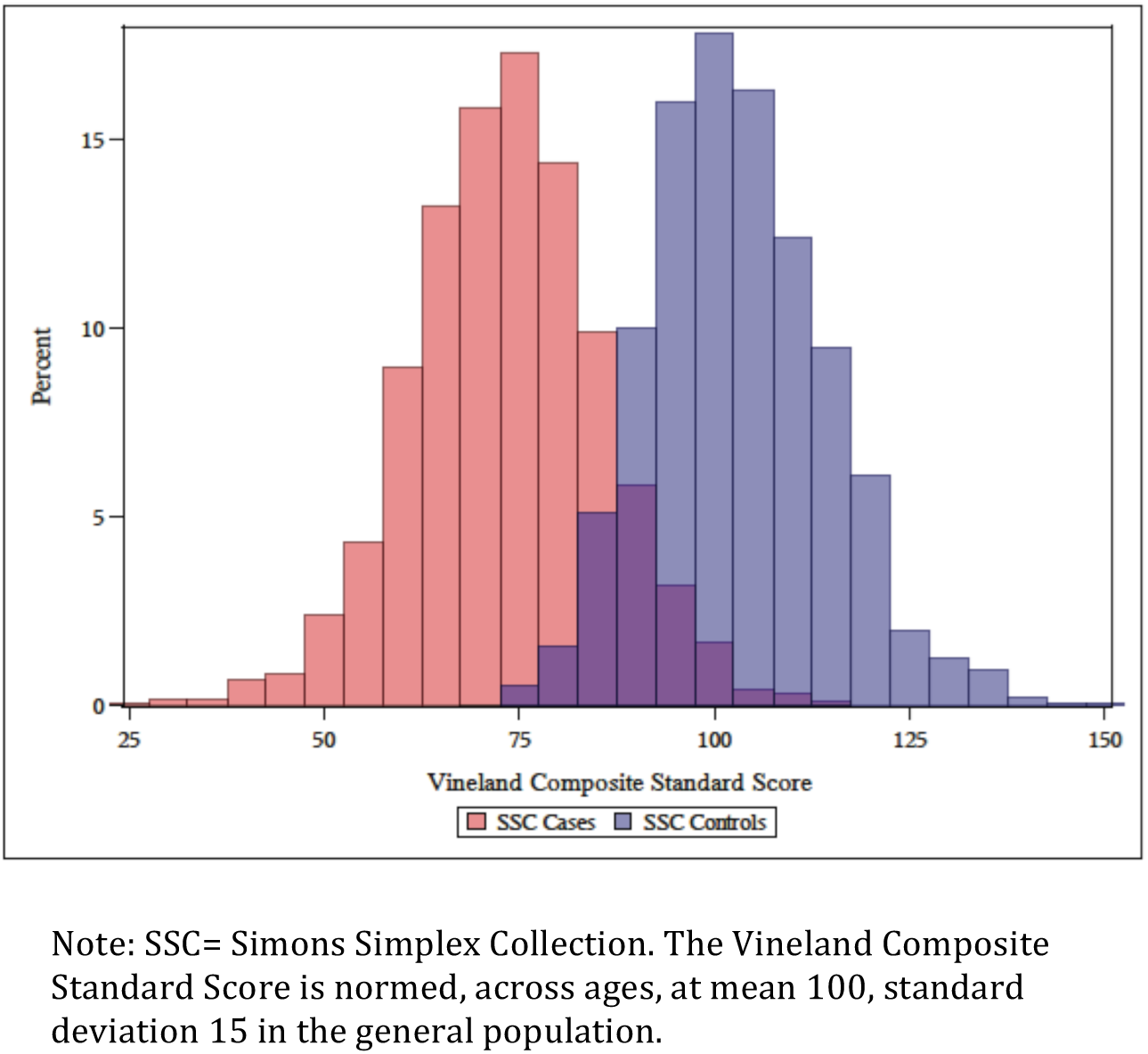
The distribution of Vineland scores overlaps between SSC cases and controls

We examined the association between Vineland scores and *de novo* variant burden in individuals with and without ASDs. Our previous work demonstrated genotype to phenotype relationships using variant classes that are strongly associated with ASD risk ^26^; subsequently these analyses focused on: 1) *de novo* loss of function (LoF) variants and 2) *de novo* missense variants predicted to be damaging by the variant annotation program PolyPhen2 and occurring in a gene known to be intolerant of heterozygous missense variation (DCM) ^30,31^. We found that variants in one of these two categories can be found in 22.1% of SSC cases and 13.2% of SSC unaffected siblings (LoF+DCM RR=1.68; p=1.8e-11; online methods). To enhance signal, we further filtered this list to remove de novo variants that were also seen in adult developmentally normal individuals in the Exome Aggregation Consortium (ExAC) resource (URLs: ExAC) ^32^. The ExAC database includes 60,706 exomes. Variants absent from that reference panel, which is a proxy for standing variation in the human population, are more likely to be deleterious. For example, 18.3% of the *de novo* LoF and DCM variants in the SSC are found in ExAC and, once those are removed, the relative *de novo* LoF+DCM burden in cases increases (LoF+DCM not in ExAC RR=1.91; p=7.6e-15).

The natural association between a) *de novo* LoF+DCM variants not seen in ExAC and b) Vineland score in SSC cases and controls is shown in Figure 2b. The LoF+DCM rate is predicted linearly by functional impairment in both cases (p=0.008) and controls (p=0.0002) using poisson regression, controlling for sex. Cases and controls with equivalent quantitative levels of functional impairment, a key component of all psychiatric diagnoses, are highly similar with regard to *de novo* variant burden, suggesting that the current categorical clinical threshold is largely arbitrary from both a phenotypic and genetic point of view. The strength of the association between LoF+DCM burden and case status (at p=7.6e-15 without controlling for Vineland scores), is only nominally significant after Vineland score is controlled for (p=0.05). The associations are weaker but similar without the ExAC filter (Figure S1).

**Figure 2b.**
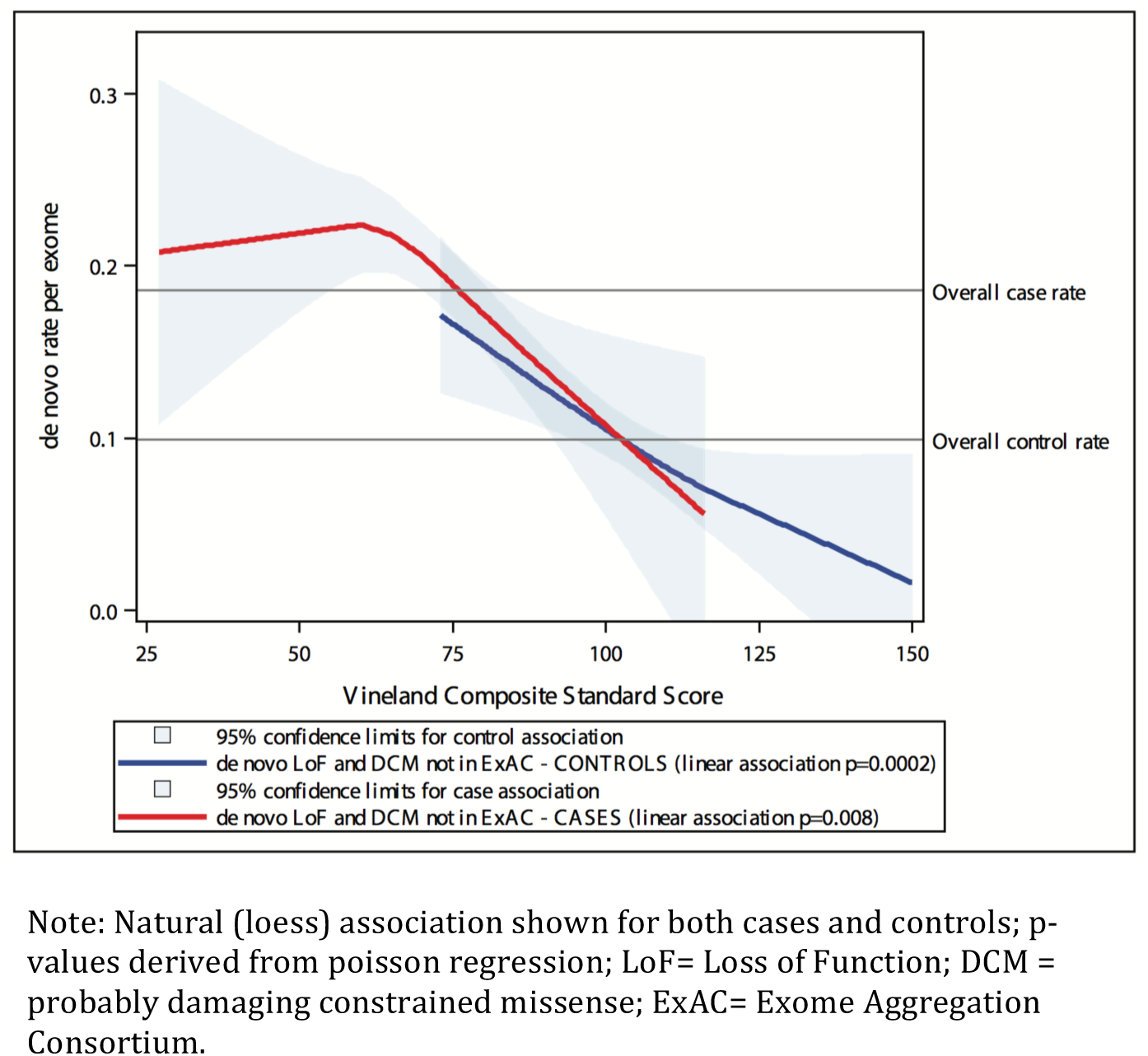
*De novo* variation influences a continuum of functional outcomes in ASD cases and controls

These data strongly suggest that genetic influences on ASD risk – both inherited and *de novo*— influence typical variation in the population in social and communication ability. They also link clinically significant problems to impairments that are less likely to be ascertained. The results have significant implications for genetic models of neuropsychiatric disorder risk. It is likely that inherited liability to ASDs is reflected in the behavioral traits of some affected individuals’ family members. This links genetic and phenotypic burden in an intuitively consistent fashion with complex, and continuously distributed, polygenic disease risk. For traits such as height, it is simple to conceptualize a model in which tall parents, (e.g. two standard deviations above the mean) are more likely to have a child that is very tall (e.g. three standard deviations above the mean). Historically this has been more complicated in neuropsychiatric disorders. Despite extensive evidence, some have even questioned the role of inheritance given that the parents of individuals with ASDs or schizophrenia rarely carry a diagnosis themselves. These results suggest that familiality should be studied in a manner beyond a count of categorically affected family members, and that trait variation in controls can provide insight into the underlying etiology of severe neurodevelopmental and psychiatric disorders. The behavioral influence of *de novo* and inherited genetic risk for ASDs can be quantified, and studies assessing continuous trait variation are likely better equipped to examine the phenotypic correlates of neuropsychiatric disease risk.

## Acknowledgements

We thank Steven Hyman, Thomas Lehner, and Nancy Kanwisher for comments on the manuscript. We are extremely grateful to all the families who took part in this study, the midwives for their help in recruiting them, and the whole ALSPAC team, which includes interviewers, computer and laboratory technicians, clerical workers, research scientists, volunteers, managers, receptionists and nurses. The UK Medical Research Council and the Wellcome Trust (Grant ref: 102215/2/13/2) and the University of Bristol provide core support for ALSPAC. Autism Speaks (7132) provided support for the analysis of autistic-trait related data in ALSPAC to BSTP. This work was also supported by the Medical Research Council Integrative Epidemiology Unit (MC_UU_12013/1-9). This publication is the work of the authors and Elise Robinson and Mark Daly will serve as guarantors for the contents of this paper. The ALSPAC GWAS data was generated by Sample Logistics and Genotyping Facilities at the Wellcome Trust Sanger Institute and LabCorp (Laboratory Corportation of America) using support from 23andMe. E.B.R. was funded by National Institutes of Mental Health Grant 1K01MH099286-01A1 and NARSAD Young Investigator grant 22379. We thank the families who took part in the Simons Simplex Collection study and the clinicians who collected data at each of the study sites. The iPSYCH project is funded by The Lundbeck Foundation and the universities and university hospitals of Aarhus and Copenhagen. Genotyping of iPSYCH and PGC samples was supported by grants from the Stanley Foundation, the Simons Foundation (SFARI 311789 to MJD), and NIMH (5U01MH094432-02 to MJD). The authors would like to thank the Exome Aggregation Consortium and the groups that provided exome variant data for comparison. A full list of contributing groups can be found at http://exac.broadinstitute.org/about.

## Authorship contributions

EBR, BSTP, VA, JK, BBS JG, KES, SS, DME, SR, MVH, TW, DMH, PBM, and ADB generated data and/or conducted analyses. EBR, BSTP, BBS, BMN, JM, DS, and MJD designed the experiment and tools. PBM, ADB, AR, GDS, and MJD supervised the research. EBR, BSTP and MJD wrote the paper.

## Methods

The datasets used in these analyses are summarized in Supplementary Table 1.

### LD Score genetic correlation analyses

Genetic correlation quantifies the extent to which two phenotypes share genetic etiology. A genetic correlation of one suggests all influences are shared; a genetic correlation of zero suggests the phenotypes are genetically independent. SNP-based genetic correlations have traditionally been estimated using data sets with individual-level genotype and phenotype data. A new method, LD Score correlation, allows one to estimate genetic relationships using SNP data when the contributing data sets are siloed ^1^. Requiring only GWAS summary statistics, the method uses linkage disequilibrium (LD) patterns to estimate genetic correlations. The resulting correlations are highly similar to those derived from residual maximum likelihood (e.g., as implemented in GCTA or BOLT-REML ^18,33^) and Haseman-Elston regression (URLs: Haseman-Elston). LD Score regression is implemented in the free and open source software package LDSC (URLs: LDSC).

We estimated the genetic association between diagnosed ASDs and traits of social and communication impairment in the general population using LD score correlation. Traits of social and communication impairment were measured using the Social and Communication Disorders Checklist (SCDC) in the Avon Longitudinal Study of Parents and Children (ALSPAC). ALSPAC is a population-based, longitudinal cohort study that initially recruited 14,541 pregnancies in Bristol, United Kingdom with expected dates of delivery between April 1, 1991 and December 31, 1992. All women in the study area with expected delivery dates in that time frame were recruited for participation ^19,20^. Ethical approval was obtained from the ALSPAC Law-and-Ethics Committee (IRB00003312) and the Local Research Ethics Committees, and written informed consent was provided by all parents. The study website contains details of all available data (URLS: ALSPAC). The GWAS of the SCDC in ALSPAC, from which we obtained the SCDC summary statistics, has already been published ^23^. Exome sequencing data from trios is not available in ALSPAC.

The SCDC is a 12-item parent-rated scale that counts traits of social and communication impairment quantitatively. Each question has a 0, 1, or 2 response option; the subsequent range of total scores is 0-24. Individuals with diagnosed ASDs, on average, have very high scores on the SCDC, consistent with the disorders’ definitional social and communication impairment ^21^. As described in the SCDC GWAS, the SCDC was used at multiple time points in ALSPAC ^23^. The SCDC is stable over time ^22^ and individual scores over time are genetically correlated ^23^. To reduce multiple testing, we used the age 8 SCDC data as childhood autistic traits are well studied, and have been linked to diagnosed ASDs through twin studies ^12,13^. We also focused on the age 8 data as it maximizes SCDC sample size (n= 5,628), and accordingly maximizes the power of the correlation tests.

The ASD case-control GWAS summary statistics come from the PGC Autism (PGC-ASD) group and the iPSYCH-SSI-Broad Autism group (iPSYCH-ASD). The PGC-ASD summary statistics were generated using publically available data from a meta-analysis of 5,305 ASD diagnosed cases of ASD and 5,305 pseudocontrols of European descent (URLS: PGC). The pseudocontrol methodology used by the PGC Autism group, and the PGC GWAS analytic pipeline, have been reported on extensively ^4,34^. Pseudocontrols are built using the untransmitted alleles from each parent at each locus. The subsequent GWAS is immune to population stratification as the pseudocontrols are ancestrally matched to the cases. LD score correlations have already been estimated using these data, as described in Bulik-Sullivan and Finucane, in press. In Figure 1, we highlight a subset of the correlations from that manuscript, specifically those associating PGC-ASD with publically available summary statistics from PGC schizophrenia, PGC bipolar disorder, and PGC major depressive disorder (URLS: PGC).

The iPSYCH-ASD data are from a new population based case-control ASD sample derived from the Danish Neonatal Screening Biobank hosted by Statens Serum Institut, comprising dried bloodspots (Guthrie cards) from all individuals born in Denmark since 1981. The samples can be linked to the Danish register system, including the Danish Psychiatric Central Register. DNA extracted from the bloodspots can be successfully amplified and employed in GWAS ^35,36^. The iPSYCH-ASD project aims to genotype all Danish individuals with an available bloodspot and an ASD diagnosis in their medical record (ICD codes F84.0, F84.1, F84.5, F84.8, and F84.9). This study has been approved by the Danish research ethical committee system. This analysis employs the iPSYCH-ASD data generated to date, specifically the first 10 genotyping waves of that collection which contain 7,783 ASD cases and 11,359 controls. All individuals in the sample were born between 1981 and 2005. Genotyping was performed at the Broad Institute. The data were cleaned and analyzed using the same analysis pipeline described in Ripke et al. 2014 and other previous PGC GWAS publications.

The GWAS summary statistics used for the secondary genetic correlation analyses are published and publicly available. Summary statistics from a GWAS of child full scale IQ have been made public by Benyamin et al. 2014, and were used in Bulik-Sullivan and Finucane et al. (in press) to estimate a genetic association between PGC-ASD and IQ in the general population. More than forty percent of contributing individuals in the Benyamin GWAS were from the ALSPAC cohort (mean age 9 years). We used LD score correlation to similarly estimate a genetic association between iPSYCH-ASD and general population full scale IQ in childhood.

Each of the genetic correlation estimates were obtained using constrained intercept cross-trait LD Score regression, which yields much lower standard errors (by ~30%) than unconstrained intercept LD Score regression ^1^. Unlike unconstrained intercept LD Score regression, constrained intercept LD Score regression can give biased estimates of genetic correlation if the two studies have either (a) hidden sample overlap or (b) shared population stratification. We can rule out both of these concerns in the present study, because the PGC-ASD sample is family-based (case-pseudocontrol), and does not have case or control overlap with any other PGC analyses. The iPSYCH-ASD and ALSPAC samples do not contain any cases or pseudocontrols that were used in PGC-ASD. In addition, case-pseudocontrol analyses are immune to confounding from population stratification, so it is not possible for genetic correlation estimates to be biased by population stratification when at least one of the studies used a case-pseudocontrol design. As a robustness check, we verified that the point estimates of genetic correlation obtained with unconstrained intercept LD Score regression are similar to the constrained intercept results presented in Figure 1 (although the standard errors are higher, due to the lower statistical efficiency of unconstrained intercept LD Score regression). These results are presented in Supplementary Table 2.

### De novo variant analyses

The Simons Simplex Collection (SSC) resource is unique in its combination of detailed phenotypic and genotypic characterizations. The SSC ascertained over 2800 individuals with ASDs and their nuclear family members, restricting recruitment to families in which no other cases of ASD have been diagnosed out to the level of first cousins. Families were also excluded in the event of intellectual disability in a sibling or a history of parental schizophrenia ^28^. To our knowledge, the SSC siblings currently constitute the most deeply phenotyped control sample with available *de novo* variant information. We used those data to examine the relationship between *de novo* variant burden and phenotypic variation in the SSC, in both siblings (n=1,861) and probands (n=2,497).

We limited the analysis to *de novo* variant classes that are strongly associated with ASD risk, specifically loss of function mutations (LoF) and probably damaging missense mutations found in constrained genes (DCM). LoF mutations include frameshift, splice site, and nonsense mutations. DCM mutations include variants that are a) predicted to be damaging by PolyPhen2 and b) occurring in a gene known to be intolerant of heterozygous missense variation ^30,31^. In the SSC, LoF mutations are found in 8.9% of controls and 15.0% of cases; DCM mutations are found in 4.3% of controls and 7.1% of cases. Restricting the classes of *de novo* variants analyzed increases the probability of association to phenotypic variation ^26^. Synonymous variants, for example, are associated with neither case status (p=0.23) nor phenotypic variation in IQ (p=0.87) or Vineland scores (p=0.91) in SSC cases. To increase signal, we further filtered observed *de novo* variants on their presence/absence in the publically available Exome Aggregation Consortium database (ExAC; URLS: ExAC). The ExAC database contains jointly called exomes from 60,706 developmentally normative adult individuals, recruited for an array of exome sequencing studies. These individual exomes form a reference panel of consistently processed human exonic variation, with particular utility towards the consideration of rare variants. Just as selection reduces the probability with which LoF variants will be seen in genes with low tolerance for functional disruption ^30^, reference genomes will be less likely to contain variants that create strong risk for reproductively deleterious phenotypes like ASDs. In other words, one expects that *de novo* variants not seen in 60,706 reference exomes will be, on average, more deleterious. We considered an SSC *de novo* variant recurrent if a variant with the same chromosome, position, reference allele, and alternate allele was seen in ExAC. After filtering out the variants seen in ExaC, the LoF+DCM rate was 19.0% in cases and 9.9% in controls.

We have previously associated *de novo* variant burden with variation in case IQ and other measures of case severity, including the Vineland Scales of Adaptive Behavior (Vineland), in the SSC ^26^. IQ was not measured in SSC siblings, however the Vineland was used in a manner consistent with its use in cases. The Vineland assesses overall adaptive functioning, with subscales assessing 1) social and 2) communication ability, as well as 3) daily living skills. The Vineland is commonly used as one measure of case severity in ASD, though its social and communication subscales are not designed to capture canonical autism-like symptomology. In this analysis, we used Poisson regression in, in both cases and controls, to associate Vineland scores with *de novo* variant burden. Sex was controlled for in all *de novo* variant analyses.

## URLs

LDSC: http://www.github.com/bulik/ldsc

Haseman-Elston: http://www.biorxiv.org/content/early/2015/04/20/018283).

ALSPAC: http://www.bris.ac.uk/alspac/researchers/data-access/data-dictionary

PGC: http://www.med.unc.edu/pgc/downloads

ExaC: http://exac.broadinstitute.org

### Collaborators

iPSYCH-SSI-Broad Collaborators: Thomas D. Als, Marie Baekvad-Hansen, Richard Belliveau, Anders D. Borglum, Mark J. Daly, Ditte Demontis, Ashley Dumont, Jacqueline Goldstein, Jonas Grauholm, Jakob Grove, Christine Hansen, Thomas F. Hansen, Mads V. Hollegaard, Daniel Howrigan, David M. Hougaard, Francesco Lescai, Julian Maller, Joanna Martin, Manuel Mattheisen, Jennifer Moran, Preben Bo Mortensen, Ole Mors, Benjamin N. Neale, Merete Nordentoft, Bent Norgaard-Pedersen, Timothy Poterba, Jesper Poulsen, Elise B. Robinson, Stephen Ripke, Christine Stevens, Raymond Walters, and Thomas Werge

## SUPPLEMENTAL TABLES AND FIGURES

**Table S1.**
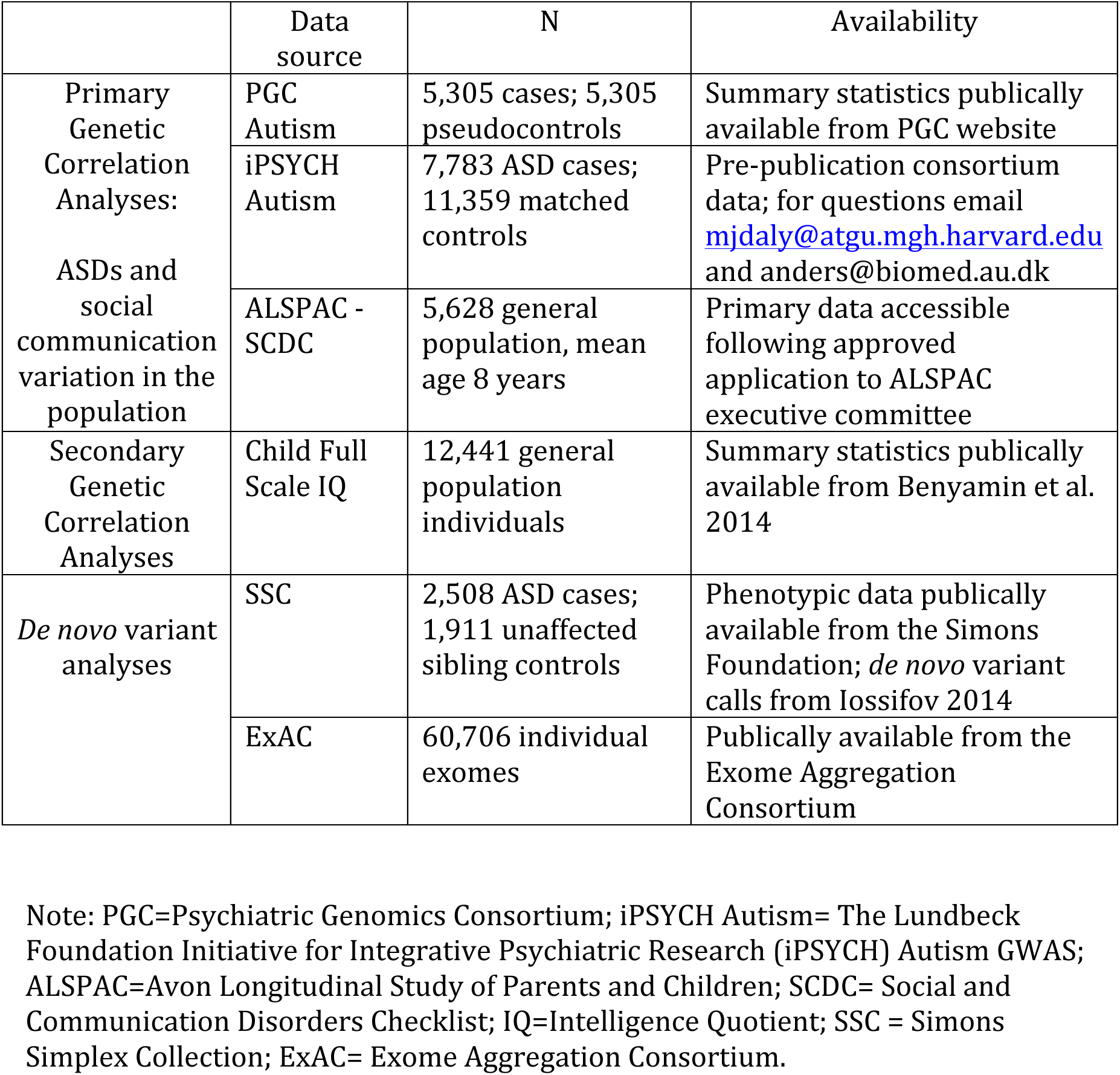
Data sources

**Table S2.**
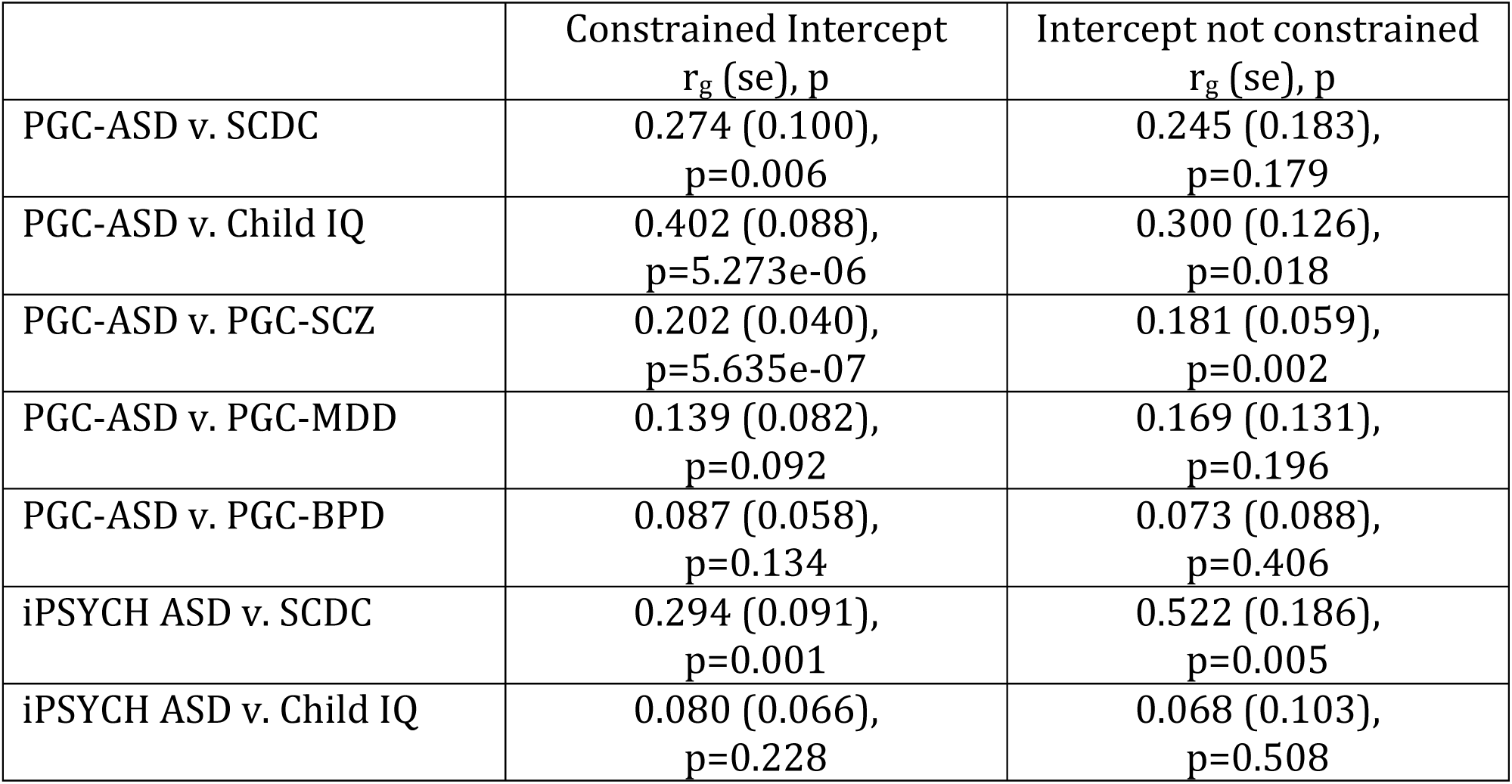
LD score genetic correlation estimates with and without intercept constraint

| | Constrained Intercept<br>$r_g$ (se), p | Intercept not constrained<br>$r_g$ (se), p |
| --- | --- | --- |
| PGC-ASD v. SCDC | 0.274 (0.100),<br>p=0.006 | 0.245 (0.183),<br>p=0.179 |
| PGC-ASD v. Child IQ | 0.402 (0.088),<br>p=5.273e-06 | 0.300 (0.126),<br>p=0.018 |
| PGC-ASD v. PGC-SCZ | 0.202 (0.040),<br>p=5.635e-07 | 0.181 (0.059),<br>p=0.002 |
| PGC-ASD v. PGC-MDD | 0.139 (0.082),<br>p=0.092 | 0.169 (0.131),<br>p=0.196 |
| PGC-ASD v. PGC-BPD | 0.087 (0.058),<br>p=0.134 | 0.073 (0.088),<br>p=0.406 |
| iPSYCH ASD v. SCDC | 0.294 (0.091),<br>p=0.001 | 0.522 (0.186),<br>p=0.005 |
| iPSYCH ASD v. Child IQ | 0.080 (0.066),<br>p=0.228 | 0.068 (0.103),<br>p=0.508 |

**Figure S1.**
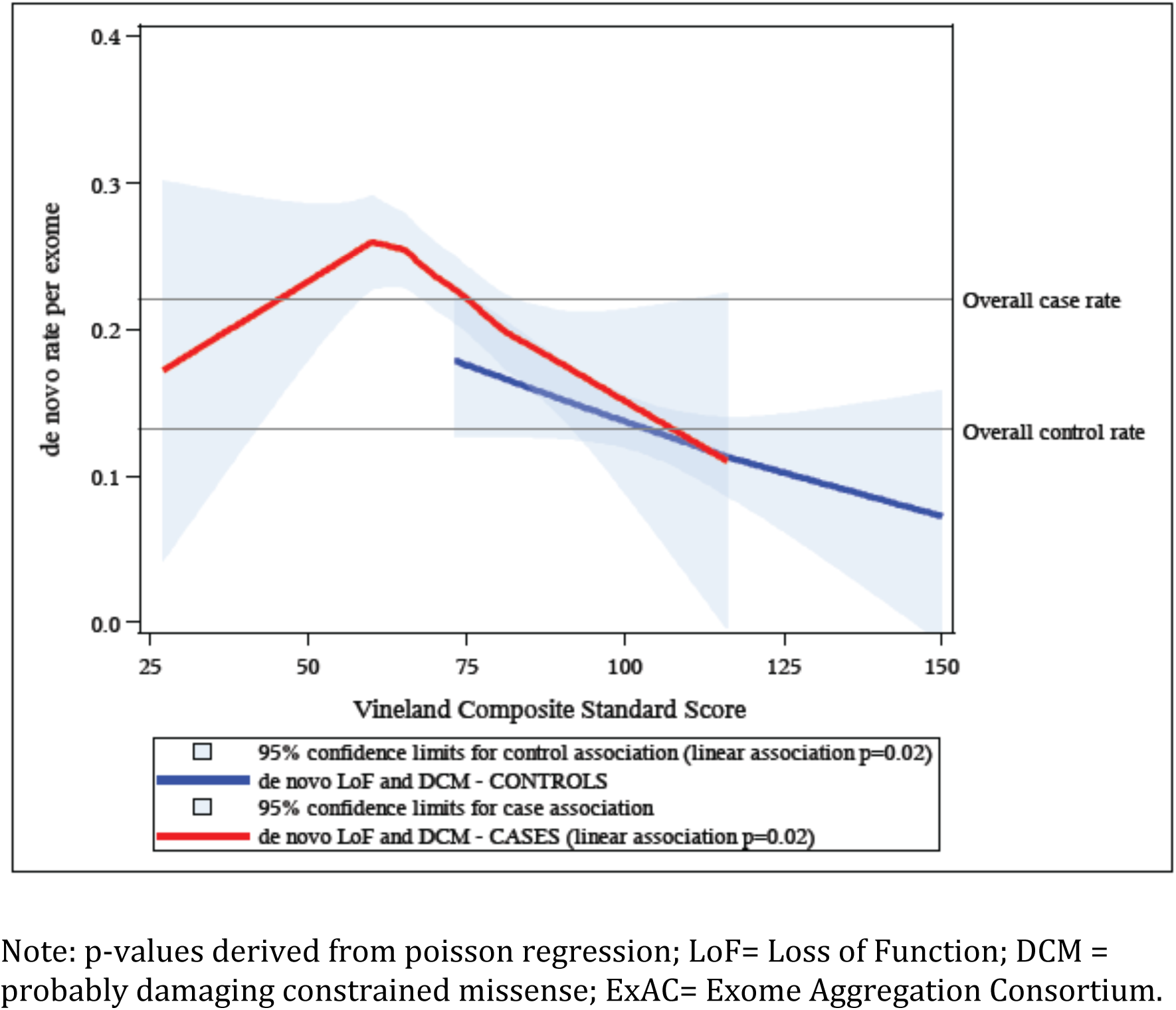
*De novo* variant continuum without the ExAC filter

